# SmProt: A Reliable Repository with Comprehensive Annotation of Small Proteins Identified from Ribosome Profiling

**DOI:** 10.1101/2021.04.29.441405

**Authors:** Yanyan Li, Honghong Zhou, Xiaomin Chen, Yu Zheng, Quan Kang, Di Hao, Lili Zhang, Tingrui Song, Huaxia Luo, Yajing Hao, Yiwen Chen, Runsheng Chen, Peng Zhang, Shunmin He

## Abstract

Small proteins specifically refer to proteins consisting of less than 100 amino acids translated from small open reading frames (sORFs), which were usually missed in previous genome annotation. The significance of small proteins has been revealed in current years, along with the discovery of their diverse functions. However, systematic annotation of small proteins is still insufficient. SmProt was specially developed to provide valuable information on small proteins for scientific community. Here we present the update of SmProt, which emphasizes reliability of translated sORFs, genetic variants in translated sORFs, disease-specific sORFs translation events or sequences, and significantly increased data volume. More components such as non-AUG translation initiation, function, and new sources are also included. SmProt incorporated 638,958 unique small proteins curated from 3,165,229 primary records, which were computationally predicted from 419 ribosome profiling (Ribo-seq) datasets and collected from the literature and other sources originating from 370 cell lines or tissues in 8 species (*Homo sapiens*, *Mus musculus*, *Rattus norvegicus*, *Drosophila melanogaster*, *Danio rerio*, *Saccharomyces cerevisiae*, *Caenorhabditis elegans*, and *Escherichia coli*). In addition, small protein families identified from human microbiomes were collected. All datasets in SmProt are free to access, and available for browse, search, and bulk downloads at http://bigdata.ibp.ac.cn/SmProt/.

## Introduction

Genome annotation is fundamental to life science. In recent years, it has been found that small open reading frames (sORFs) widely exist in genomes of many organisms including human [1] and human microbiomes [2], and some are able to be translated into small proteins [3–5]. Small proteins are proteins with less than 100 amino acids, which may derive from untranslated regions (UTRs) of mRNAs [6] or non-coding RNAs [7,8] including pri-miRNAs [9,10], lncRNAs [11], and circRNAs [12]. Small proteins were usually missed in previous coding sequence annotation, while their significance has been revealed in current years for diverse functions [13], such as embryonic development [14,15], cell apoptosis [16], muscle contraction [17], and antimicrobial activity [18]. Some are proved to play roles in many diseases [19,20] including tumors [9,11,12]. Despite the abundance of sORFs in genome, the number of well-studied small proteins is very limited. Annotation of numerous small proteins will contribute to studies on various physiology and pathology processes.

Identification of small proteins at proteomic level is challenging. Mass spectrum (MS) can provide direct evidence of small proteins, but it relies much on the coverage of existing libraries, which mainly focused on large proteins rather than small proteins. Protease cleavage sites were lacking in small proteins limited by length. Besides, small proteins are usually of low abundance, and tend to be filtered out during enrichment process [21]. Ribosome profiling (also named Ribosomal footprinting or Ribo-Seq) provides a more sensitive way for global detection of translation events based on the deep sequencing of ribosome-protected mRNA fragments (RPFs) [22,23], which allows for identifying the location of translated ORFs and translation initiation sites, the distribution of ribosomes on mRNA, and the speed of translating ribosomes [24]. Reference library for mass spectrometry can also be constructed with Ribo-Seq results. The regular Ribo-seq (rRibo-seq) utilizes cycloheximide (CHX) [25], a drug binding at the ribosome E-site [26], as a translation elongation inhibitor to freeze translating ribosomes. Translation is principally regulated at the initiation stage. TI-seq is a variation of rRibo-seq technique that use different translation inhibitors, usually lactomidomycin (LTM) [25] or harringtonine (HAR) [27], which can induce ribosomes stasis at translation initiation (TI) sites (TISs). TI-seq enables the global mapping of translation initiation sites, and is more accurate in prediction of non-AUG start codons. Many sORFs are proved to use non-classical AUG start codon [28], which is also an important mechanism for generating protein isoforms [29,30]. rRibo-seq data usually have clear triplet periodicity [26]. Different computational analysis strategies [31–38] have been developed to identify translated sequences using Ribo-seq data.

Emerging evidences showed that many upstream open reading frames (uORFs) act in cis to regulate the translation of downstream ORFs by leaky scanning [39], reinitiation [40], and ribosome stalling [41]. Recently, variants creating new upstream start codons or disrupting stop sites of existing uORFs (uORF-perturbing) were found under strong negative selection [42]. uORF-perturbing variants were demonstrated as an under-recognized functional class that contribute to human disease.

Since great importance has been attached to small proteins, in-depth investigations of small proteins across various species are in need. SmProt is dedicated to integrate knowledge of proteins shorter than 100 amino acids (hereinafter referred to as small proteins) translated from various sources, especially for ones from UTRs and non-coding RNAs. The annotation information and functional sections in the current release are much richer than those in the 1^st^ release [43], and the data volume and reliability are greatly improved.

## Data collection and processing

### Data sources

rRibo-seq and TI-seq datasets derived from diverse tissues/cell lines were collected from GEO database [44] and European Nucleotide Archive [45]. The latest reference genomes and gene annotation were download from Ensembl [46], GENCODE [47], and NCBI-Genome database. WGS Variants were collected from their respective websites. The construction pipeline of SmProt was summarized as follows (**Figure 1**).

**Figure 1.**
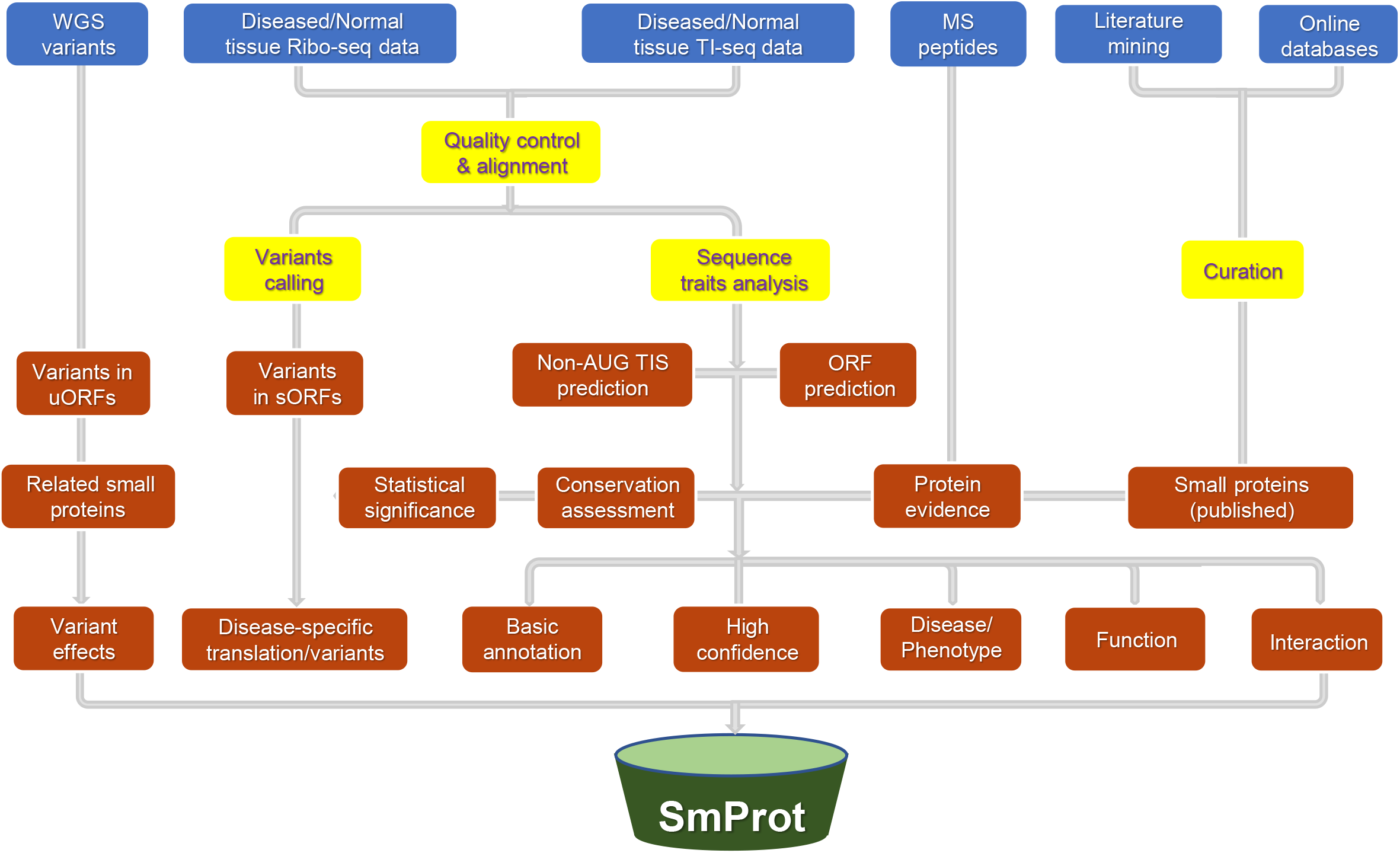
Construction pipeline of SmProt. Blue background: data sources. Yellow background: management processes. Red background: results. Abbreviations: WGS, whole genome sequencing; MS, mass spectrometry; TIS, translation initiation site; ORF, open reading frame; sORF, small open reading frame; uORF, upstream open reading frame.

### Ribo-seq data processing

The fastq files of 547 Ribo-seq datasets were downloaded from GEO and European Nucleotide Archive database. Each dataset was checked manually to confirm the sequencing adapters. The adapters were removed using cutadapt 1.18 [48] and only reads with length 25–35 were kept. Then the sequences were mapped to the latest genome using STAR 2.5.2a [49] using EndToEnd mode with allowance of up to 2 mismatches.

Ribo-seq quality and P-site offsets were assessed by Ribo-TISH [34] quality module. For TI-seq data, more attention was put on TIS quality (-t). Manual checks were then carried out to verify offset values and eliminate datasets without obvious triplet periodicity. After the quality control, 419 Ribo-seq datasets (Supplementary Table S1) were kept.

Translated ORFs were predicted by Ribo-TISH predict module. Biological and technical duplication data under the same treatment in one dataset were merged. Minimum amino acid length of candidate ORF was set to 5. Considering both ATG and near-cognate start codons, rRibo-seq datasets using only CHX without matched TI-seq were analyzed twice. One is prediction of ORFs with canonical ATG start codon, the other is prediction of ORFs with near-cognate start codons with 1 base different from ATG (--alt). Preferring data evidence instead of prior assumption in our database, only the best frame test results from multiple candidate start codons in the same ORF were reported (--framebest). For datasets containing TI-seq data, alternative start codons were included (--alt), and different parameters were set for LTM-based TI-seq and HARR-based TI-seq (--harr).

sORFs with less than 100 amino acids were filtered from the above prediction results. To avoid confusion from classic proteins, those marked as *known* (means the translation initiation site is annotated in another transcript)*, CDSFrameOverlap* (means the ORF overlaps with annotated CDS in another transcript in the same reading frame), and *Truncated* (means the ORF is part of annotated CDS in the same transcript) without translation initiation evidence (i.e. none significant results identified from paired TI-seq datasets) were further removed, considering these results may be supported by RPFs from other classic proteins longer than 100 AAs.

In-frame reads of sORFs were counted and normalized by library sequencing depth (in-frame total reads count) and sORF length, a similar method with RPKM (Reads Per Kilobase per Million mapped reads) in RNA-seq but using ribosome profiling data which represents the translation levels.

Finally, 3,060,793 records were kept. Results with the identical genome loci in one species were merged as the same small protein generating 577,206 unique IDs, while information derived from multiple datasets were kept, a similar integration method to piRBase [50].

### Variants from ribosome profiling data

We performed germline variants detection on 96 human ribosome profiling datasets, referring to the workflow for processing RNA data for germline short variant discovery with GATK v4.1.8 [51–54]. Duplicate reads were identified using MarkDuplicates tool after alignment, then reads with N in Cigar were split using SplitNCigarReads tool. Base quality score recalibration was carried out based on true sites in training sets using BaseRecalibrator tool and applied using ApplyBQSR tool. Variants were called individually in each sample using the HaplotypeCaller tool. Variants with QualByDepth (QD) < 2 were removed using VariantFiltration tool. Germline single nucleotide variants (SNVs) were linked to small proteins in SmProt according to genomic positions.

### Variants from WGS data

Variants from 1KGP3 [55], GAsP [56], TOPMed [57], gnomAD3 [42,58], and NyuWa [59] were collected. VCF files were lifted over from old genome version to GRCh38 using LiftoverVcf tool of GATK with allowance to recover swapped ref and alt alleles.

Variants in 5’UTRs were evaluated for their effects on translated uORFs in SmProt using VEP [60] with plugin UTRannotator [42,61], and classified by their functional consequences.

### Disease-specific small proteins

Small proteins identified only from diseased cell lines/tissues but not from corresponding normal cell lines/tissues were predicted as disease-specific translation events: if there were matched data of normal and diseased groups in the same dataset, small proteins derived uniquely from diseased group were screened as disease-specific ones; if there’s no matched control group in the same dataset, the same type of healthy tissue/cell line in other datasets were used as control. If there’s no matched same tissue/cell line, all data from diverse normal tissues/cell lines were merged for comparisons (Supplementary Table S2), and small proteins identified only from the diseased cell lines/tissues were predicted as tissue-specific. Disease-specific or tissue-specific translation events require RiboPvalue in disease groups under 0.01 while similar proteins with different TISs at the same loci in control group not detected (RiboPvalue higher than 0.05).

Single nucleotide variants (SNVs) detected only in diseased variant sets but not in normal sets were predicted as disease-specific SNVs. SNVs in diseased cell lines/tissues derived from ribosome profiling data and located within the genomic region of small proteins were regarded as diseased variant sets. SNVs in corresponding normal cell lines/tissues (Supplementary Table S2) derived from ribosome profiling data were combined with all variants derived from multiple WGS projects, as control variant sets for comparison.

### Function domain prediction

Besides function of small proteins collected from literature mining, we used InterProScan [62] to predict function domain of small proteins, which focuses on combination of protein family membership and the functional domains/sites, and has been extensively used by genome sequencing projects and the UniProt Knowledgebase [63]. Default thresholds and additional parameters -*goterms -pa* were adopted for gene oncology and pathway annotations.

### PhyloCSF calculation

Pre-calculated BigWig data of PhyloCSF [64] scores at each base across the whole genome were downloaded from the broad institute https://data.broadinstitute.org/compbio1/PhyloCSFtracks/, and the score for genomic region of each small protein was extracted with our script using pyBigWig (https://github.com/deeptools/pyBigWig).

### Database implementation

Database website was organized with HTML (https://html.spec.whatwg.org/), JavaScript (https://www.javascript.com/), PHP (https://www.php.net/), and MYSQL (https://www.mysql.com/). UCSC Genome Browser (http://genome.ucsc.edu/) was used to visualize the small proteins and variants. NCBI BLAST (https://blast.ncbi.nlm.nih.gov/Blast.cgi) was used for sequence similarity searches.

## Database content and usage

### Overview

SmProt was constructed by pipeline described in Figure 1. Multiple ways were provided to search, browse, visualize, and study small proteins (**Figure 2**). Small proteins were found mainly from rRibo-seq and TI-seq data. All information for small proteins from different data sources and datasets were integrated. General information for small proteins was provided such as sequence, mass, location, blocks, tissue or cell line, predicted functions, conservation, and multiple IDs including small protein ID, Ensembl ID, and NONCODE [65] ID. Translation level (in frame counts and Ribo RPKM) of small proteins identified from each dataset and record was provided. Details for their related variants and diseases were also provided (**Figure 3**). SmProt now has 638,958 unique small proteins and 3,165,229 small protein records in total (**Table 1;** Supplementary Table S3).

**Figure 2.**
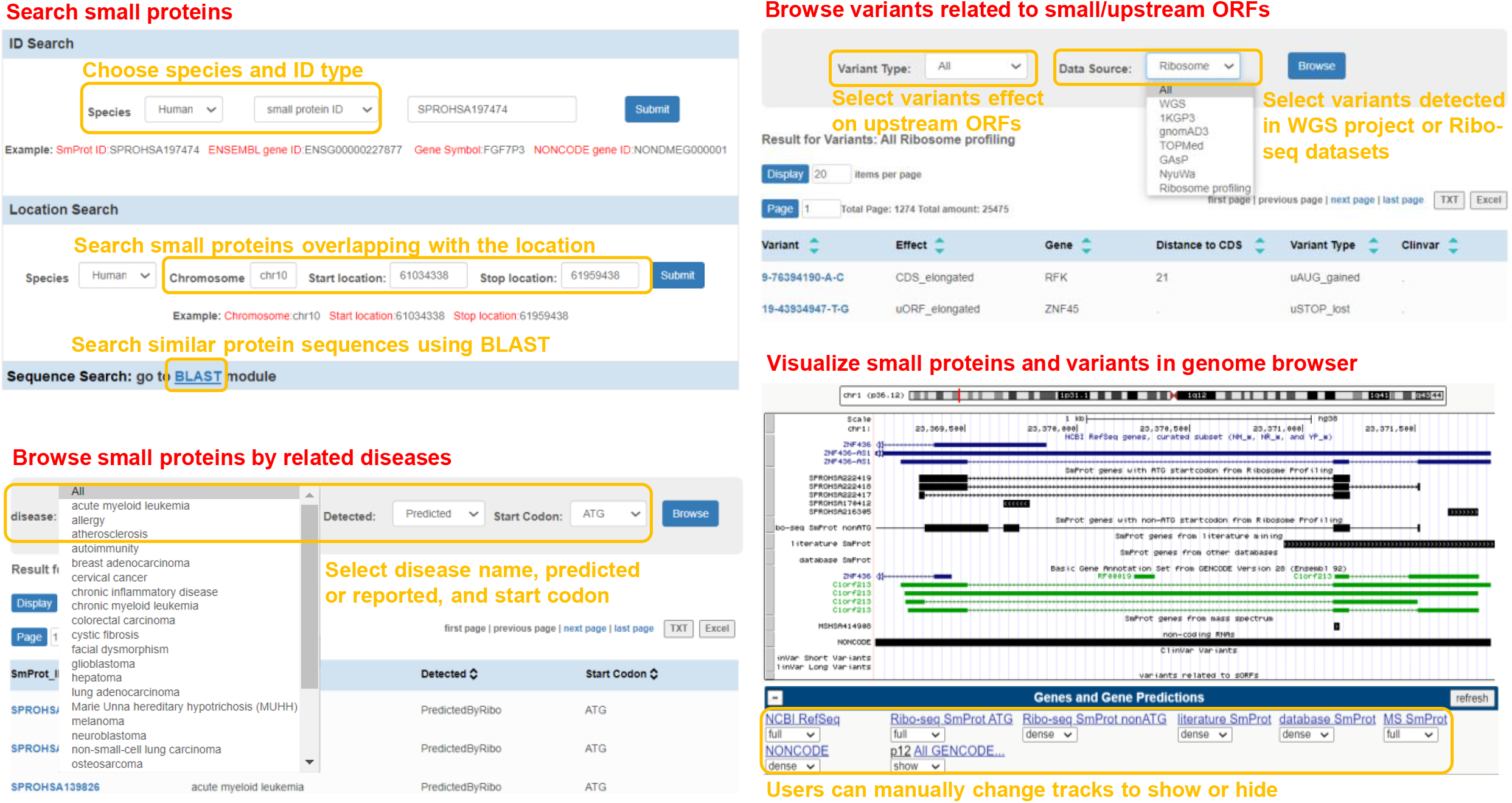
Usage of SmProt. SmProt provided multiple ways to search, browse, visualize small proteins, related diseases, and variants. Abbreviations: WGS, whole genome sequencing; ORF, open reading frame.

**Figure 3.**
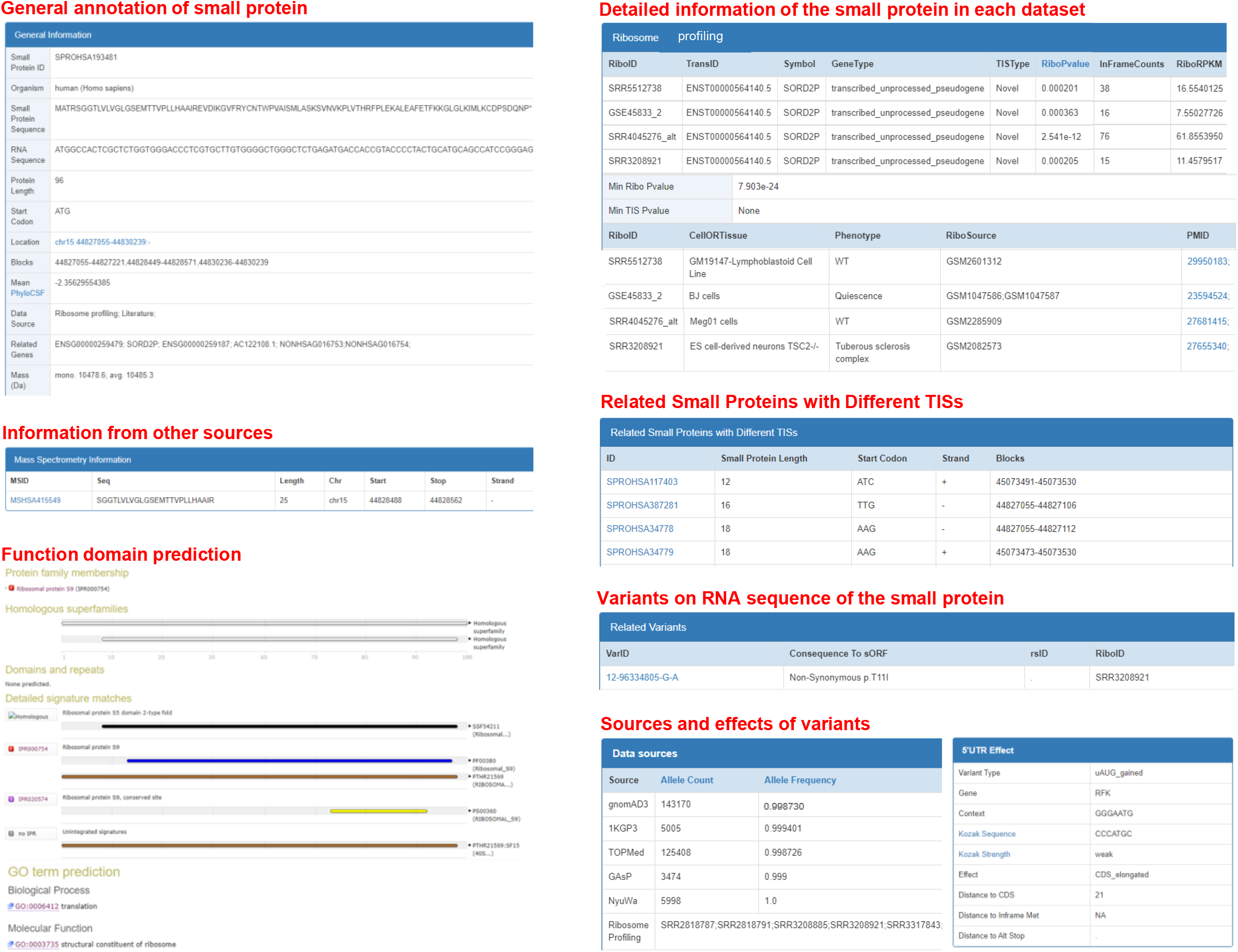
Contents of SmProt. Detailed information for small proteins, including general annotation, information from ribosome profiling data, literature, other databases, mass spectrometry, function domain prediction, related diseases, related variants from WGS projects as well as corresponding effects, etc. Abbreviations: WGS, whole genome sequencing; TIS, translation initiation site.

**Table 1.** Statistics of unique small proteins in SmProt.

| Type | Start codon | Human | Mouse | Fruit fly | Rat | C.elegans | Yeast | E.coli | Zebrafish | All |
| --- | --- | --- | --- | --- | --- | --- | --- | --- | --- | --- |
| Ribo-seq | ATG | 70,931 | 48,909 | 5269 | 3560 | 4334 | 4535 | 1881 | 1924 | 141,343 |
|  | Near ATG | 229,653 | 133,037 | 29,679 | 9910 | 9894 | 12,339 | 10,004 | 1347 | 435,863 |
| Literature | All | 38,157 | 8875 | 22,228 | 163 | 4 | 355 | 296 | 3612 | 73,690 |
| Databases | All | 786 | 797 | 100 | 271 | 120 | 336 | 955 | 64 | 3429 |
| MS | All | 768 | 51 | 66 | 38 | 0 | 3 | 0 | 1 | 927 |
| All IDs | All | 327,995 | 189,433 | 56,574 | 13,829 | 14,255 | 17,312 | 12,881 | 6679 | 638,958 |

*Note:* Not including human microbiomes small protein families. IDs mean unique entries with identical genome loci in one species. Abbreviations: Ribo-seq, ribosome profiling; MS, mass spectrometry.

### Reliability of small proteins

SmProt emphasizes reliability of small proteins, which is guaranteed mainly by the significance of 3 nt periodicity in RPF P-site profile:

Firstly, we constructed new pipeline based on independently published toolkit Ribo-TISH [34], which allows for accurate detection of ORFs and TISs using rRibo-seq and TI-seq. Ribo-TISH uses rank sum test to detect 3 nt periodicity, and negative binomial test to detect translation initiation sites, which outperforms other established methods in prediction accuracy.

Secondly, in addition to the quality control based on Ribo-TISH quality module, manual checks were also carried out to ensure clear triplet periodicity and unambiguous offset of Ribo-seq data, which further eliminated noises.

Thirdly, we provided several evaluations as evidences: p-values of small proteins called from multiple ribosome profiling datasets indicating the confidence in different samples and conditions; PhyloCSF conservation of genomic regions reflecting coding potential; and peptide evidence derived from mass spectrum data. All evidences were exhibited in the small protein page. What’s more, a set with evidence of both translation events and protein fragments was provided on download page.

What’s more, information of small protein derived from multiple sources were integrated in small protein information page.

### Variants related to small proteins

25,475 variants located on translated sORFs were provided, which were exhibited in the related small protein page. For uORF-perturbing variants are likely to impact translation of downstream proteins [42], variants from multiple WGS projects and ribosome profiling data were evaluated for their effects on translated uORFs in SmProt, which can be found at variants page.

### Disease-specific small proteins

Disease-specific small proteins have potential to be candidates of molecular markers or targets for diagnosis and treatment. Disease-specific translation events as well as disease-specific SNVs of small proteins in 16 types of diseases were identified (see methods) (Supplementary Table S4). Besides, small proteins that have been verified experimentally in certain diseases were also documented through literature mining.

### Human microbiomes small proteins

Over 4000 conserved small protein families identified from human microbiomes were collected [2]. A new section *HumanMicroBio* was created to integrate and display selected information of these small protein families.

### Other sources

We use a set of keywords (Supplementary File S1) to search articles about small proteins in PubMed database. High-confidence small proteins in CCDS [66] and Swiss-Prot [67] were also integrated. Literature mining is processed in stages, and the newly published data from other sources is being released continuously after accomplishment of manual review and curation.

### Function domain prediction

For successfully predicted functions of small proteins derived from ribosome profiling and literature mining, SmProt provided graph for visualization and prediction details including gene oncology (GO) and pathway annotation. Users can choose *predicted functions* on *Browse* page to filter the results with function domain prediction.

### Inner BLAST

The abundant small proteins across multiple species allows for sequence similarity searches of both nucleotides and proteins. Users can search for sequences of interests using BLASTp and BLASTx (NCBI BLAST 2.2.24 release) online.

### Visualization using UCSC Genome Browser

SmProt incorporated UCSC Genome Browser [68] for visualization of all the information including genomic loci of small proteins, variants from ribosome profiling data and multiple WGS projects related to small proteins, MS data, and gene annotation. The latest genome versions including hg38, mm10, rn6, dm6, ce11, sacCer3, and danRer11 were provided.

### Comparison with other databases

SmProt currently includes 419 Ribo-seq datasets derived from 116 cell lines/tissues, compared to 60 datasets derived from 37 cell lines/tissues in the initial version. The number of small protein records identified from ribosome profiling in the current release is 60 times that of the 1^st^ release (3 million to 0.05 million). The current release of SmProt combined a large amount of duplicate records in 1^st^ release [43], and Ribo-seq analysis pipeline was optimized to ensure the reliability of our results. Variants in translated sORFs identified from Ribo-seq data as well as uORF-perturbing variants identified from WGS projects were provided. Disease-specific small proteins may provide new perspectives for clinical studies.

Currently, there are a few databases for small proteins such as ARA-PEPs [69], PsORF [70], and sORFs.org [71]. ARA-PEPs and PsORF only harbors small proteins in plants. sORFs.org developed simple inner TIS-calling algorithm not based on triplet periodicity, which should be the most important feature of Ribo-seq. SmProt emphasizes high confidence using our Ribo-TISH pipeline that is more accurate than previous methods. SmProt analyzed 419 Ribo-seq datasets, while there were only 78 in sORFs.org. SmProt pays special attention to function, variants, and related diseases of small proteins, and WGS data resource are also integrated, which other databases didn’t pay attention to.

Other proteomic databases such as UniProt, neXtProt [72], and OpenProt [73] are not specifically designed for small proteins. neXtProt only harbors proteins of human while SmProt harbors small proteins in 8 species. OpenProt also used ribosome profiling and mass spectrum to predict proteins including some small proteins longer than 30 amino acids, while SmProt analyzed much more ribosome profiling datasets (419), which were about 5 times that in OpenProt (87), and provided small proteins longer than 5 amino acids.

## Conclusion

In brief, SmProt integrated small proteins from large amount of ribosome profiling data, and provides more abundant details. We strongly believe that SmProt will provide valuable and accurate information on small proteins for scientific community. Moreover, it provides a new resource for users interested in function and mechanism study, and a reference for construction of mass spectrometry library of small proteins.

## Data Availability

SmProt is publicly available at http://bigdata.ibp.ac.cn/SmProt/.

## CRediT author statement

**Yanyan Li:** Conceptualization, Methodology, Investigation, Formal analysis, Data Curation, Writing - Original Draft, Software, Visualization. **Honghong Zhou:** Investigation, Data Curation, Funding acquisition. **Xiaomin Chen:** Investigation, Data Curation. **Yu Zheng:** Data Curation, Software, Visualization. **Quan Kang:** Software, Visualization. **Di Hao:** Data Curation, Software. **Lili Zhang:** Visualization. **Tingrui Song:** Visualization. **Huaxia Luo:** Writing - Review & Editing. **Yajing Hao:** Writing - Review & Editing. **Yiwen Chen:** Software. **Runsheng Chen:** Resources, Supervision, Funding acquisition. **Peng Zhang:** Conceptualization, Methodology, Investigation, Software, Writing - Review & Editing, Visualization, Project administration, Funding acquisition. **Shunmin He:** Conceptualization, Methodology, Resources, Investigation, Writing - Review & Editing, Supervision, Funding acquisition.

## Acknowledgments

This work was supported by National Natural Science Foundation of China (Grant Nos. 81902519, 31871294, 31701117, and 31970647); the National Key R&D Program of China (Grant Nos. 2016YFC0901702 and 2018YFA0106901); the Strategic Priority Research Program of Chinese Academy of Sciences (Grant No. XDB38040300); the 13th Five-year Informatization Plan of Chinese Academy of Sciences (Grant No. XXH13505-05); special investigation on science and technology basic resources of the MOST, China (Grant No. 2019FY100102); the National Genomics Data Center. We thank Center for Big Data Research in Health (http://bigdata.ibp.ac.cn/), Institute of Biophysics, Chinese Academy of Sciences, for supporting data analysis and computing resource.

## Supplementary material

**Supplementary Table S1: Information of ribosome profiling datasets analyzed in SmProt (.xlsx)**

**Supplementary Table S2: Dataset contrasts for generating disease-specific small proteins and variants (.xlsx)**

**Supplementary Table S3: Statistics of small proteins primary records in SmProt (.xlsx)**

**Supplementary Table S4: Statistics of small proteins and variants specific in diverse diseases (.xlsx)**

**Supplementary File S1: Keywords for literature mining (.doc)**

