## Supplementary File S1 for "SmProt: A Reliable Repository with Comprehensive Annotation of Small Proteins Identified from Ribosome Profiling"

**#Keywords for literature mining**

“small protein”, “small peptide”, smORF, sORF, sPEP, “small open reading frame”, “small ORF”, “short open reading frame”, “short ORF”, micropeptide, miPEP, miRNA & ORF, miRNA & “open reading frame”, “peptide-encoding ORF”, “coding small ORF”, peptide & ORF, “decoding sORF”

“downstream ORF”, “upstream ORF”, uAUG, uORF, dORF, uPEP, upORF, “upstream AUG”, “upstream initiation”, “upstream open reading frame”, “downstream open reading frame”,

ORF&UTR, “upstream start codon”, “upstream translation initiation”, UTR & peptide

non-AUG & sORF, “non-canonical initiation codon” & sORF, non-AUG & “open reading frame”, “non-canonical initiation codon” & “open reading frame”, non-AUG & peptide, non-AUG start codon, “repeat-associated non-AUG” & peptide, “non-canonical initiation codon” & peptide,

non-canonical initiation codon,

ribo-seq & peptide, “ribosome footprints” & peptide, “ribosome profiling” & peptide, “ribosome protection” & peptide

ribo-seq & ORF, “ribosome footprints” & ORF, “ribosome profiling” & ORF, “ribosome protection” & ORF

ribo-seq & “open reading frame”, “ribosome footprints” & “open reading frame”, “ribosome profiling” & “open reading frame”, “ribosome protection” & “open reading frame”

TI-seq, QTI-seq, Ribo-RET, polyribosomal RNA-seq, MS

circRNA & sORF, “decoding circRNA”, noncoding & peptide, sORF & lncRNA, circRNA & peptide, “circular RNA” & peptide, pri-miRNA & ORF
